# COVATOR: A Software for Chimeric Coronavirus Identification

**DOI:** 10.1101/2020.11.14.383075

**Authors:** Peter T. Habib

**Author notes:** **Corresponding Author:** Peter T. Habib, Bioinformatics Specialist, Colors Medical Laboratories.

## Abstract

The term chimeric virus was not popular in the last decades. Recently, according to current sequencing efforts in discovering COVID-19 Secrets, the generated information assumed the presence of 6 Coronavirus main strains, but coronavirus diverges into hundreds of sub-strains. the bottleneck is the mutation rate. With two mutation/month, humanity will meet a new sub-strain every month. Tracking new sequenced viruses is urgently needed because of the pathogenic effect of the new substrains. here we introduce COVATOR, A user-friendly and python-based software that identifies viral chimerism. COVATOR aligns input genome and protein that has no known source, against genomes and protein with known source, then gives the user a graphical summary.

## Introduction

The largest known single-stranded RNA viruses are coronaviruses (CoVs) [1]. They cause a different illness in mammals and birds, ranging from enteritis in cows, pigs, and upper respirational diseases in chickens to life-threatening humans. The first diagnosed case in Southern China was a severe acute respiratory syndrome (SARS) that was first described in 2002 [2]. Then just 10 years after the appearance of a SARS-CoV, a new evolving Coronavirus called Middle East Respiratory Syndrome (MERS-CoV) affected about 50% of people in the Middle East. on 10 March 2016, The Global MERS Case Count according to the World Health Organization (WHO) was 1,651 laboratory-confirmed cases [3]. The murin coronavirus of the SARS and MER-SCoV viruses, Mouse Hepatitis Virus (MHV), was a model for both the molecular biology and the pathogenesis of the members of these viral families for many years.

Quite recently, a novel beta CoV coronavirus (2019 nCoV) has been connected to serious human respiratory infections from the province of Wuhan, China. In China, new 8 cases of 2019 nCoV related pneumonia, plus approximately 650,000 cases from 23 other countries, have been reported at the time of the written study. Now, this pathogen has 1,301,818 deaths (Source: WorldMeter). Phylogenetic relationships between Bat and Human coronaviridae have been discovered for SARS [4] and more recently also for 2019□nCoV,[5] suggesting events of inten□species transmissions.[6]

Viruses, such as SARS-CoV-2, HIV, and influenza, encode their genome in RNA, tend to collect mutations as copied within their hosts, since enzymes that copy RNA are susceptible to errors. In the latest days, sequencing evidence indicates that coronaviruses transform slower than most other RNA viruses, presumably because of an enzyme that reads and remedies lethal copying errors. Just two mutations per month in virus genome accumulate representing about half the influenza rate and a quarter HIV rate [7]. Most modifications do not modify the actions of the infection, but a few can alter the transmissibility or seriousness of the illness.

Korber et al. demonstrated in April 2020 that the “D614G” transition had contributed to the rising transmissibility of viruses that soon became the world’s dominant SARS-CoV-2 lineage [8]. The variant, however, could make the vaccines easier to target SARS-CoV-2. However, extreme acute respiratory syndrome (SARS) started to circulate in humans, a deletion mutation that might have hindered its propagation was created. [9]

More time, more mutations, more different strains. For example, one mutation in its receptor-binding domain (RBD) was found to be in India out of two isolates at location 407 and it was not previously recorded. At that place, isoleucine (a hydrophobic amino acid which is also a C-β branched amino acid, has been substituted for arginine. [10]

At the beginning of the pandemic, we were dealing with only one version of the virus. Now, we are dealing with virus lineage-specific for each country, soon we are going to deal with the unique virus for each group of people, and the worst scenario when it will be a unique version for each person. Tracking the virus through countries is urgently needed. There are many software tracks changes in virus genome such as Nextstrain [11] and genome-protein classification such as COVIDier [12] but it is still limited to submitted data only. Here we introduce COVATOR, a real-time tracking software where the user could submit a virus genome and the results will show the locations where protein sequences are originated.

## Material and Methods

The idea behind COVATOR is using BLAST [13] to align genome against genome and proteins with known source locations. It firstly aligns genome against genome to get an overview of the genome source, secondly align referenced proteins against the inputted genome to get a deeper insight into each protein source.

### Database Construction

COVATOR uses “makeblastdb” software to build two reference databases, one for genome and one for proteins. reference genomes to align inputted unknown genome against reference genomes with known location to find out how unknown inputted genome like references and the second is reference CDs to discover the source location of each protein to get more details about proteins source of the unknown inputted genome.

### BLASTing

Using “blast”, COVATOR begins alignment of all coronavirus twelve CDs against the inputted genome. Alignment results were formatted, sorted, and aligned in a separate file for each gene. We used CDs of the coronavirus gene to avoid protein sequence conservation and avoid the nucleotide noise in raw gene sequence where CDs have the requirement for protein-coding and each CDs has its own signature that allows the differentiation between several country sources.

### Representing the Results

COVATOR generates two folders, “CDsResults” and “GenomesResults”. Each folder contains a folder titled with the name of the inputted genome. In “GenomesResults”, BLAST aligns each CDs, python script generates a file with a summary of blast results and generates a summary figure of the country. In “CDsResults”, four Alignment files are generated. Aliview [13] format, BLAST results in FASTA format, HTML format, and BLAST result in text format.

## Discussion

The evolving virus leads to the spreading of different strains in the world, each country begins to develop its specific strain. Developing different strains may lead to confusion in drug and vaccine development. Due to mutation that may occur in critical sites in protein that represent a drug target, drug or vaccine that is effective with certain strains may not work with another strain due to mutation effect that allows viruses to disguise from treatment [10]. This diversity within the same virus leads to the production of “Chimeric” viruses in which their components are derived from different sources which are explained in figure (1).

**Figure 1:**
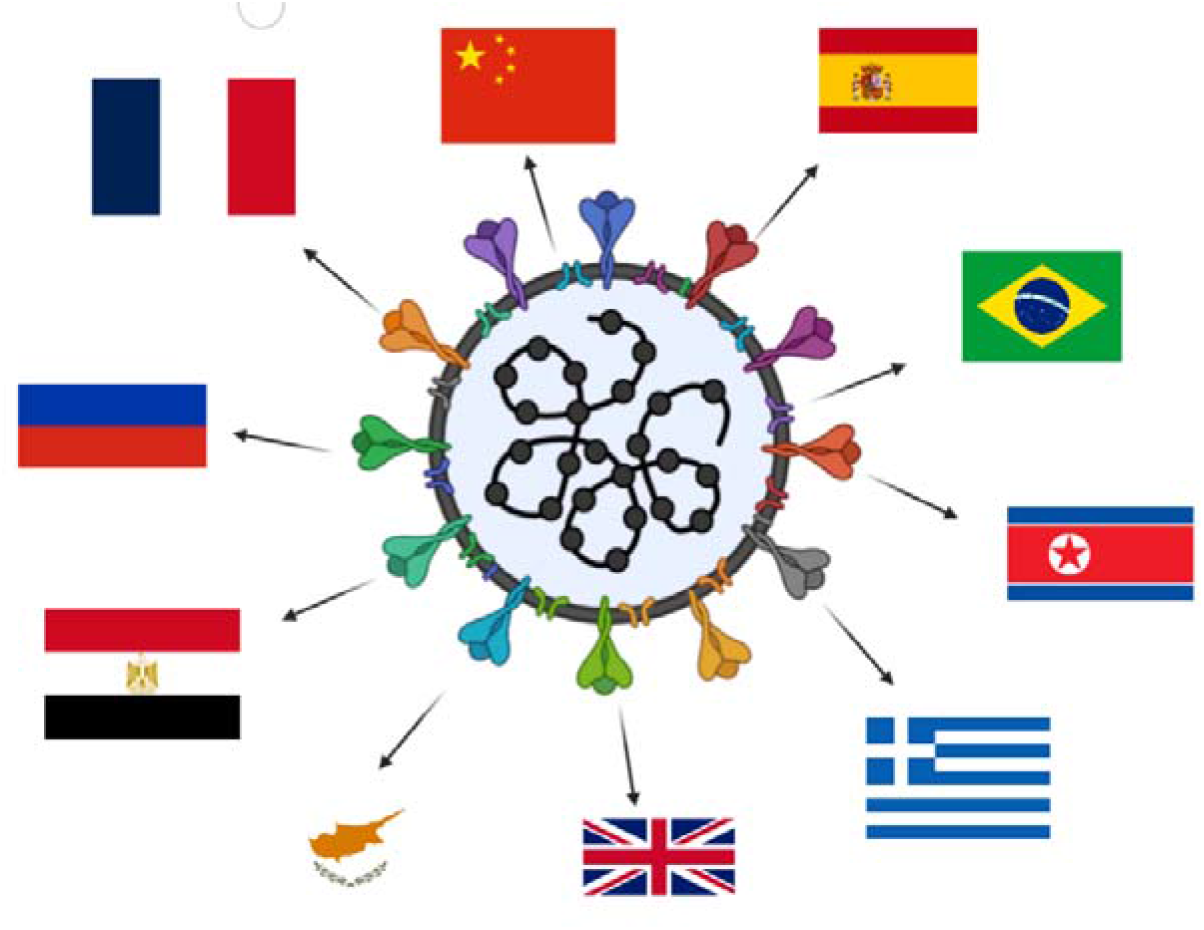
The Chimeric virus generated from cross-infection with different strain may lead to the development of new strains in which components derived from different countries

COVATOR aligning the unknown input genome with known genomes to finds out the most similar genome to the inputted one. Inputted genome passes with two steps. First is general alignment to get a fast summary of the genome source. The second, identifying each protein source by extracting unknown protein from the inputted genome, then aligning it against proteins with known source as shown in figure (2).

**Figure 2:**
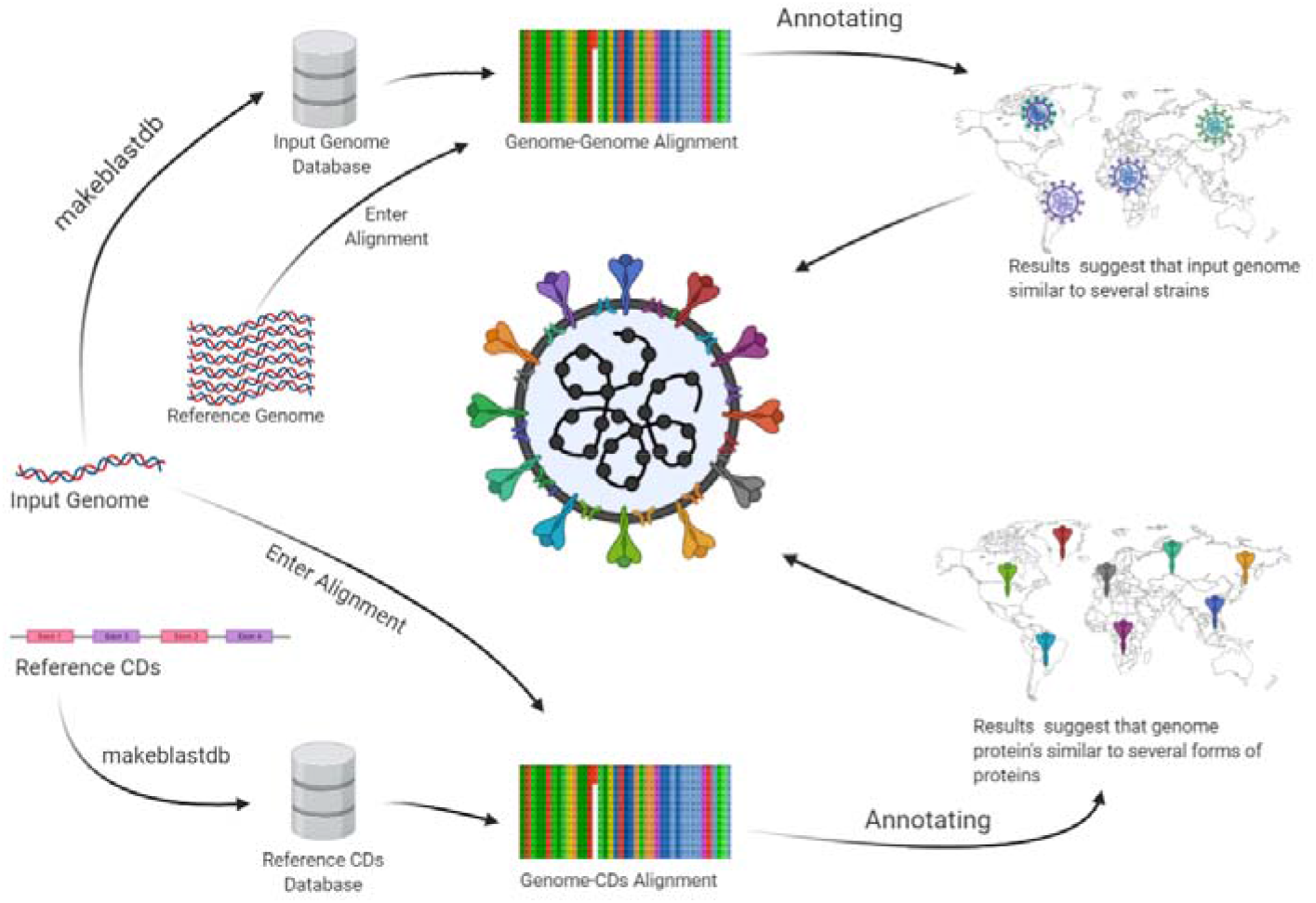
Flow chart of COVATOR

### Use Case

COVATOR analysis is composed of two steps, the first step is a summary, and the second extract deep information. We used Coronavirus genome ID: MT883499.1 from NCBI [14] in the FASTA file in the Linux Terminal (figure 3). and the results were a summary list of BLAST results (Table 1) and a Figure represent the country occurrence (figure 4). COVATOR creates a database from the input genome then blasting all CDs against it.

**Figure 3:**
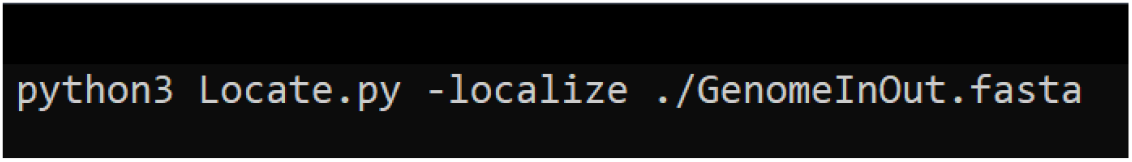
A Terminal screenshot of COVATOR command line used in running analysis

**Figure 4:**
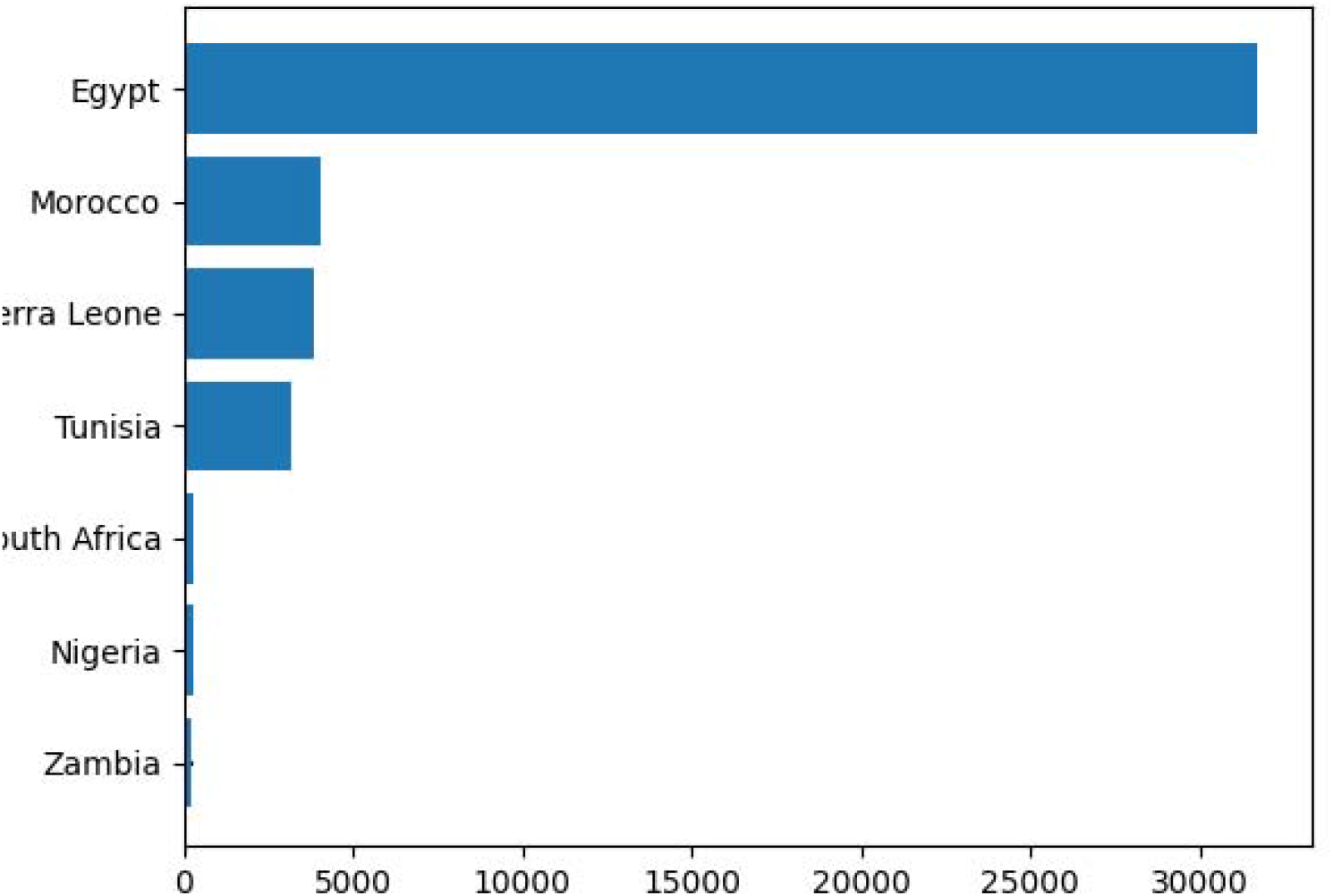
histogram shows the occurrence of each country that is has similarity hits with inputted genome

**Table 1:** Example of Generated Results from Step. Columns order: CD ID, protein name, country, inputted genome ID, and identity percent.

|  |  |  |  |  |
| --- | --- | --- | --- | --- |
| MT511065.1:26185..26412 | envelope protein | Egypt | MT883499.1 | 100 |
| MT511065.1:26185..26412 | envelope protein | Egypt | MT883499.1 | 100 |
| MT511065.1:26185..26412 | envelope protein | Egypt | MT883499.1 | 100 |
| MT511065.1:26185..26412 | envelope protein | Egypt | MT883499.1 | 100 |
| MT511065.1:26185..26412 | envelope protein | Egypt | MT883499.1 | 100 |
| MT511065.1:26185..26412 | envelope protein | Tunisia | MT883499.1 | 100 |
| MT511065.1:26185..26412 | envelope protein | Morocco | MT883499.1 | 100 |
| MT511065.1:26185..26412 | envelope protein | Morocco | MT883499.1 | 100 |
| MT511065.1:26185..26412 | envelope protein | Morocco | MT883499.1 | 100 |
| MT511065.1:26185..26412 | envelope protein | Morocco | MT883499.1 | 100 |
| MT511065.1:26185..26412 | envelope protein | Egypt | MT883499.1 | 100 |
| MT511065.1:26185..26412 | envelope protein | Egypt | MT883499.1 | 100 |
| MT511065.1:26185..26412 | envelope protein | Sierra Leone | MT883499.1 | 100 |
| MT511065.1:26185..26412 | envelope protein | Sierra Leone | MT883499.1 | 100 |
| MT511065.1:26185..26412 | envelope protein | Egypt | MT883499.1 | 100 |

The second step gets more detail about the differences between the inputted genome and reference CDs. In this step, COVATOR built a database from CDs instead of the input genome to generate the result in the targeted format needed for further analysis. BLAST results processed using view [12] to generate HTML file (figure 5), Clustal [16] alignment in FASTA format (figure 6), raw BLAST result in format 0 (figure 7), and Alignment format suitable for AliView software (figure 8)

**Figure 5:**
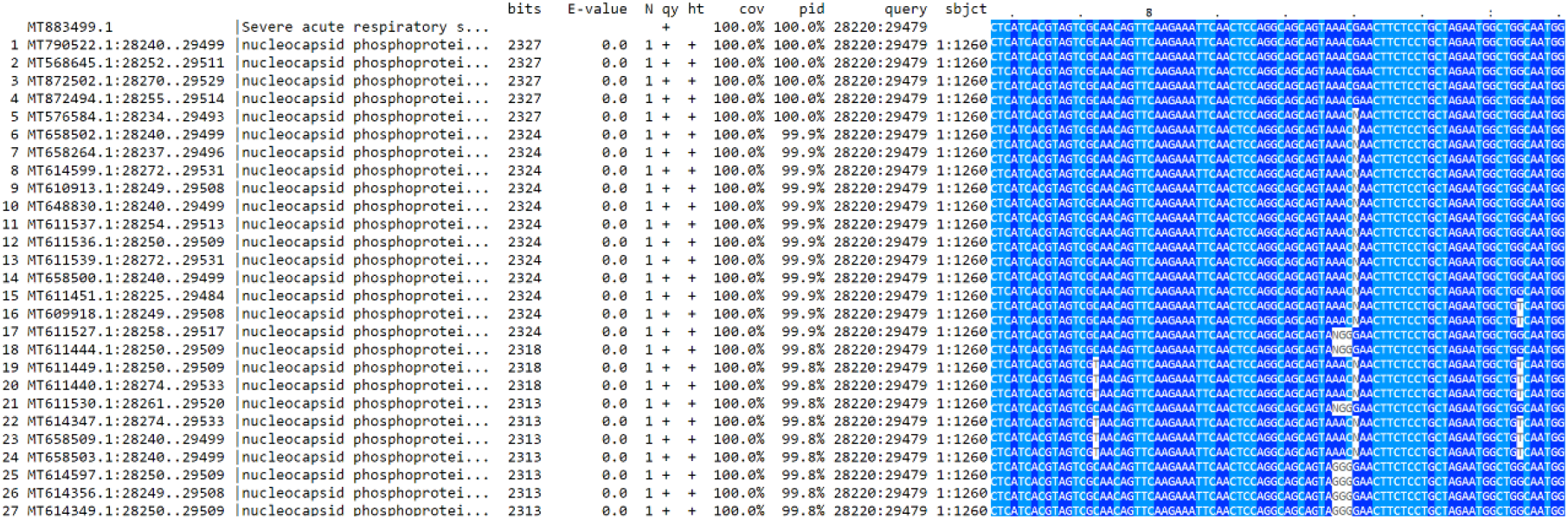
HTML view of alignment shows the differences between different countries version of a single protein which is nucleocapsid phosphoprotein

**Figure 6:**
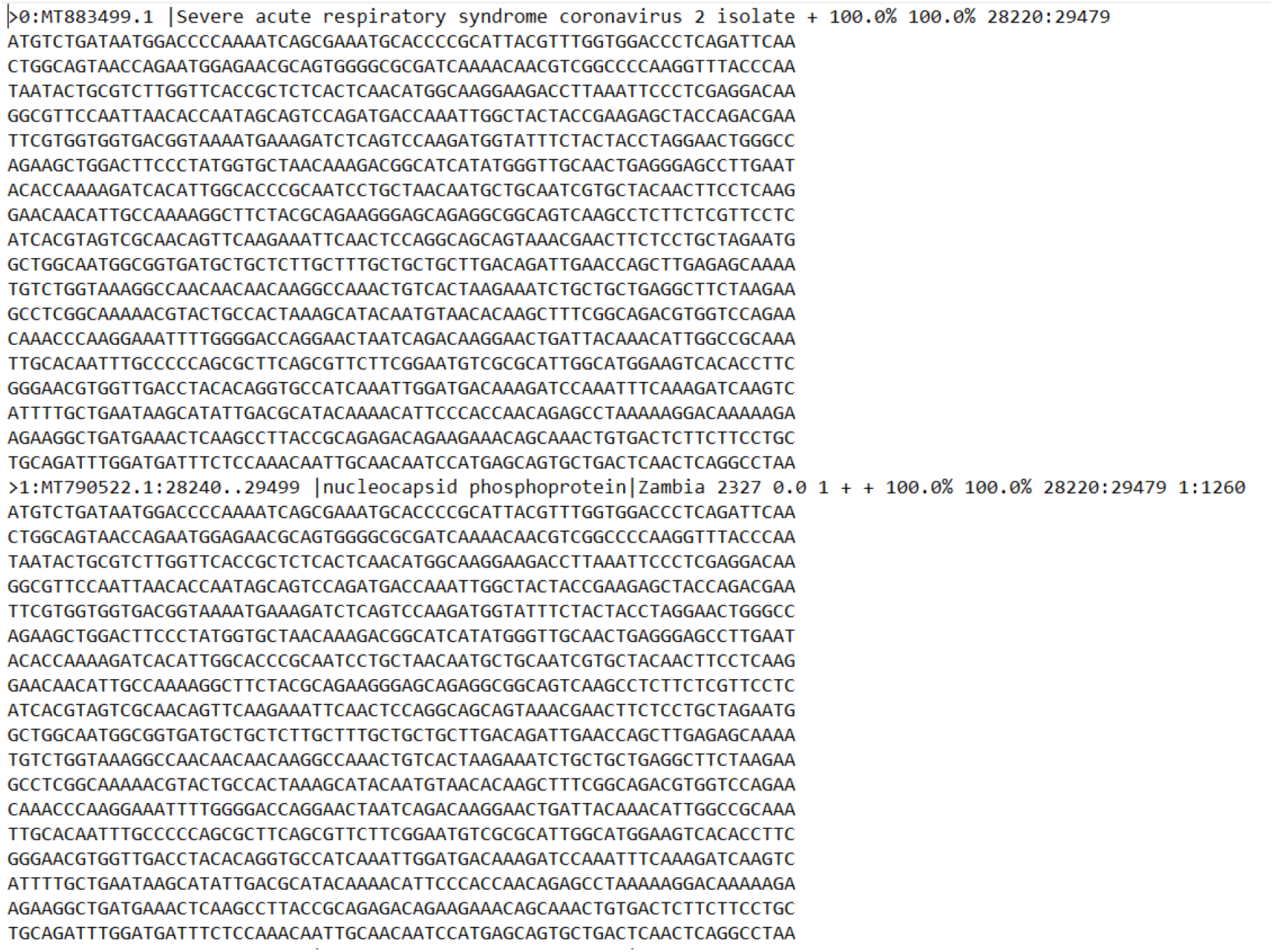
Multiple Sequence Alignment (MSA) from Clustal in FASTA format

**Figure 7:**
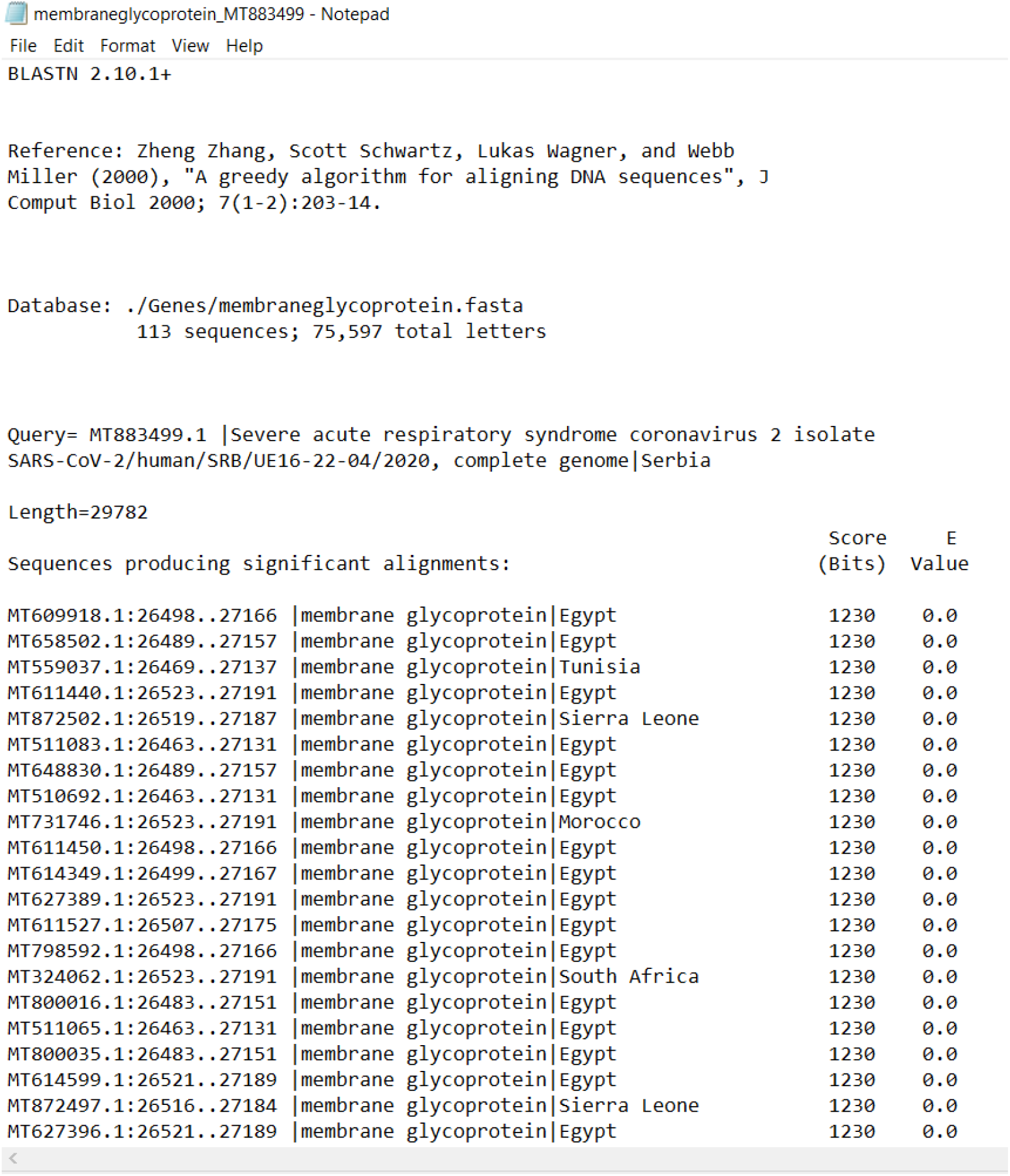
Raw BLAST output which converted using mview [17] to different formats

**Figure 8:**
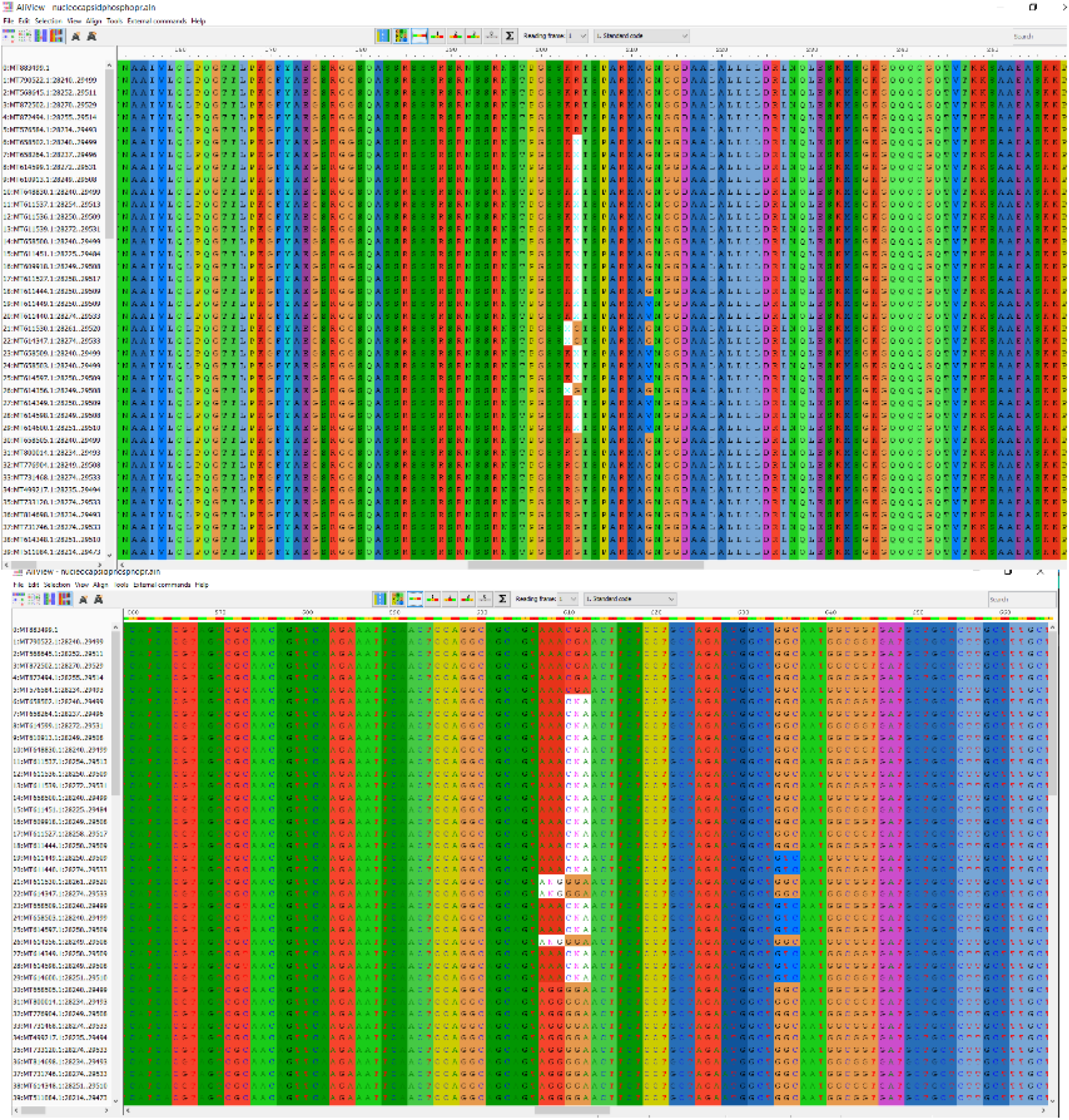
AliView Software screen shows differences in nucleotide codons (Down) and its effect on amino acid substitution (Top)

COVATOR uses Plotly [18] and webweb [19] python libraries to visualize the distribution of results. This option can be used by the command line for network map construction (figure 9) and connection map between countries in which similar protein might be found (figure 10) and both results open in an ordinary HTML browser.

**Figure 7:**
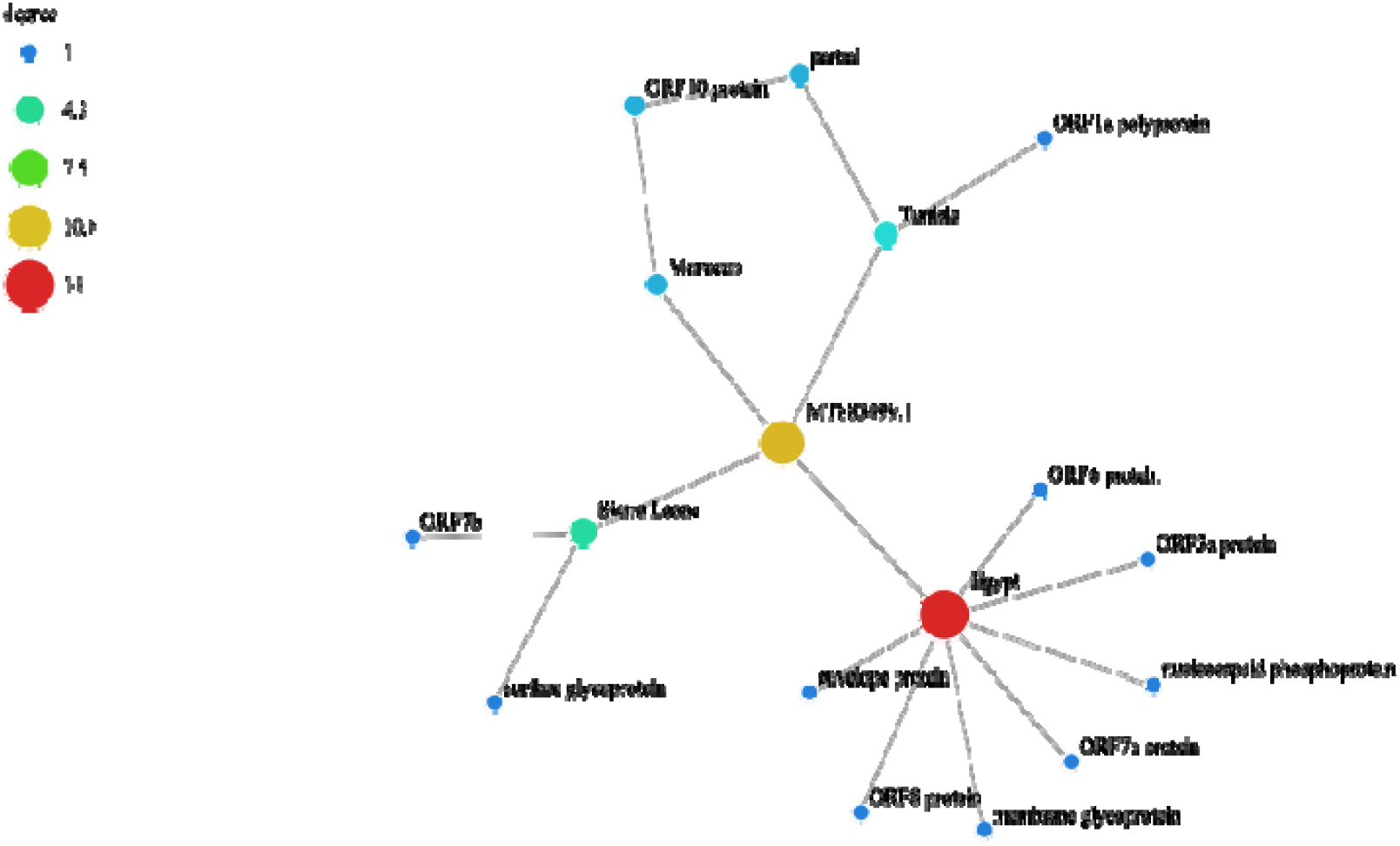
Network connection shows the input genome (yellow circle) and other countries and their proteins similar to input genome protein

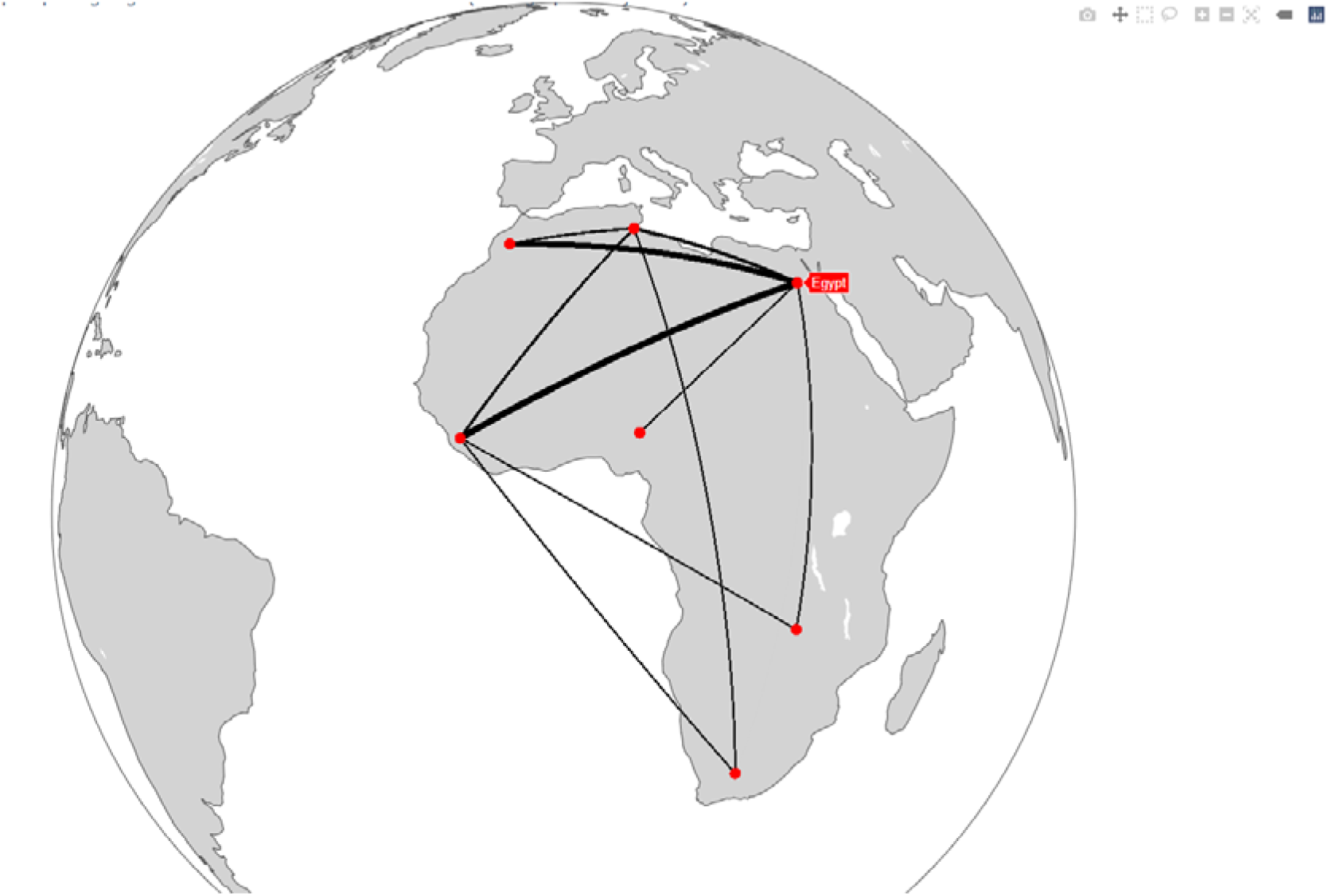

### Conclusion

The current viral bluster represents one of the biggest human health threats. The molecular and computational effort to face this killer is urgently needed. Developing new drugs or vaccine Running in full swing. Unfortunately, the virus evolved in a faster rhythm leading to emerging of new strains. Currently, we have six strains over the world according to Nextstrain [13], but the long-lasting virus in each country and mixing of different strains lead to the development of new Chimeric viruses which their proteins are sourced from different locations. The efforts in developing antiviral agent require specific details for the target of the antiviral may go in vain because of the mutations that occur within every two months. Detecting new Chimeric viruses will be the main concern of scientists soon. Here we introduce COVATOR, A new software for detecting Chimeric viruses by scanning the virus genome and determine the main sources of the protein by aligning the input unknown genome against. COVATOR is user-friendly and python 3+ based software could be running through only command line from Linux terminal.

## Data Availability

Example results and Main Scripts Available freely on Zenodo: 10.5281/zenodo.4274250

## References

1. Cui J, Li F, Shi Z□L. Origin and evolution of pathogenic coronaviruses. Nat Rev Microbiol. 2019;17(3):181□192. https://doi.org/10.1038/s4157□01□011□9

2. Du L, He Y, Zhou Y, Liu S, Zheng BJ, Jiang S. The spike protein of SARS-CoV-a target for vaccine and therapeutic development. Nat Rev Microbiol. 2009; 7: 226–236. https://goo.gl/T2kyhu

3. World Health Organization. Middle East respiratory syndrome coronavirus (MERS-CoV) updates. 2012 September 23 to 2016; January 6. 2015: https://goo.gl/NAfKqv

4. Hu B, Ge X, Wang L□F, Shi Z. Bat origin of human coronaviruses. Virol J. 2015;12(1):221. https://doi.org/10.1186/s1298□01□042□1

5. Benvenuto D, Giovannetti M, Ciccozzi A, Spoto S, Angeletti S, Ciccozzi M. The 2019□new coronavirus epidemic: evidence for virus evolution. J Med Virol. 2020. https://doi.org/10.1002/jmv.25688

6. Lu H, Stratton CW, Tang Y. Outbreak of pneumonia of unknown etiology in Wuhan China: the mystery and the miracle. J Med Virol. 2020. https://doi.org/10.1002/jmv.25678

7. Kupferschmidt, K. (2020). The pandemic virus is slowly mutating. But does it matter?

8. Korber, B., Fischer, W., Gnanakaran, S. G., Yoon, H., Theiler, J., Abfalterer, W., … & Partridge, D. G. (2020). Spike mutation pipeline reveals the emergence of a more transmissible form of SARS-CoV-2. bioRxiv.

9. Muth, D., Corman, V. M., Roth, H., Binger, T., Dijkman, R., Gottula, L. T., … & Toplak, I. (2018). Attenuation of replication by a 29 nucleotide deletion in SARS-coronavirus acquired during the early stages of human-to-human transmission. Scientific reports, 8(1), 1–11.

10. Saha, P., Banerjee, A. K., Tripathi, P. P., Srivastava, A. K., & Ray, U. (2020). A virus that has gone viral: amino acid mutation in S protein of Indian isolate of Coronavirus COVID-19 might impact receptor binding, and thus, infectivity. Bioscience Reports, 40(5).

11. Hadfield, J., Megill, C., Bell, S. M., Huddleston, J., Potter, B., Callender, C., … & Neher, R. A. (2018). Nextstrain: real-time tracking of pathogen evolution. Bioinformatics, 34(23), 4121–4123.

12. Habib, P., Alsamman, A. M., Saber-Ayad, M., Hassanein, S. E., & Hamwieh, A. (2020). COVIDier: A Deep-learning Tool For Coronaviruses Genome And Virulence Proteins Classification. bioRxiv.

13. Altschul, S. F., Gish, W., Miller, W., Myers, E. W., & Lipman, D. J. (1990). Basic local alignment search tool. Journal of molecular biology, 215(3), 403–410.

14. Larsson, A. (2014). AliView: a fast and lightweight alignment viewer and editor for large datasets. Bioinformatics, 30(22), 3276–3278.

15. Brister, J. R., Ako-Adjei, D., Bao, Y., & Blinkova, O. (2015). NCBI viral genomes resource. Nucleic acids research, 43(D1), D571–D577.

16. Larkin, M. A., Blackshields, G., Brown, N. P., Chenna, R., McGettigan, P. A., McWilliam, H., … & Thompson, J. D. (2007). Clustal W and Clustal X version 2.0. bioinformatics, 23(21), 2947–2948.

17. Brown, N. P., Leroy, C., & Sander, C. (1998). MView: a web-compatible database search or multiple alignment viewer. Bioinformatics (Oxford, England), 14(4), 380–381.

18. Sievert, C., Parmer, C., Hocking, T., Chamberlain, S., Ram, K., Corvellec, M., & Despouy, P. (2017). plotly: Create Interactive Web Graphics via ‘plotly. js’. R package version, 4(1), 110.

19. Wapman, K. H., & Larremore, D. B. (2019). webweb: a tool for creating, displaying, and sharing interactive network visualizations on the web. Journal of Open Source Software, 4(40), 1458.

